# Mechanotransduction and Biophysical Regulation of Fibroblast Plasticity Under Prolonged Shear Stress

**DOI:** 10.1101/2025.03.03.641342

**Authors:** Neha Paddillaya, Anahita Jayaram, Paturu Kondaiah, Namrata Gundiah

## Abstract

Complex interactions between mechanical stimuli and biochemical signaling shape the evolving fibrotic milieu. Fibroblasts contribute to this process, transitioning into myofibroblasts on stiffer substrates, and under low shear in 3D environments. The time-dependent effects of prolonged physiological shear stress on fibroblasts however remain poorly understood. We demonstrate that sustained shear stress (1.2 Pa) over 12 hours induces a temporal cascade of signaling and biophysical changes, driving fibroblast trans-differentiation into myofibroblasts. Cells adopt a spindle-shaped morphology under shear, actively remodeling through actin and α-SMA to form elongated stress fibers. Myofibroblasts exhibit increased vinculin and zyxin expression in mature adhesions that remain elevated post-shear. Traction stresses progressively increased over the shear duration and persisted for 12 hours after the removal of the stimulus, indicating mechanical memory. Post-shear, deadhesion strengths increased significantly, accompanied by shifts in gene expressions governing extracellular matrix homeostasis. Prolonged shear enhanced MSD, directionality, and migration speed, all crucial for effective tissue repair. Finally, we demonstrate temporal changes in the biochemical gene expressions mediating the shear-induced remodeling of the extracellular matrix, cytoskeleton, and focal adhesions during the myofibroblast transition. These findings provide insights into the mechanobiological factors driving fibroblast-to-myofibroblast transitions which may inform the development of mechano-therapeutic strategies.

## INTRODUCTION

Fibrotic scar formation is driven by the accumulation and activation of fibroblasts into large fusiform-shaped myofibroblasts through a process of fibrogenesis^1,2^. Fibroblasts are highly migratory following activation, demonstrate upregulation of proliferative and cytokine markers, and increased production of extracellular matrix (ECM) proteins and matrix metalloproteinases (MMPs)^3^. Synergistic interactions between the mechanical stimuli and biochemical signaling are highly relevant in fibrosis due to ischemic cardiomyocyte death, lung injury, cancers, or wound healing. For example, infiltrating fibroblasts in an ovine aneurysm model of the myocardial tissue replaces injured cardiomyocytes with randomly oriented collagen, disrupting electrical activity, leading to ventricular arrhythmias (Figure 1A, 1B)^4,5^. The injured tissue subsequently undergoes significant remodeling due to collagen crosslinking and tissue compaction during the proliferative stage of fibrosis, resulting in increased stiffness^6–9^. Higher tissue stiffness, a hallmark of fibrotic and chronic inflammatory disease, further activates quiescent fibroblasts into myofibroblasts. This activation alters cellular mechanotransduction, reinforcing and amplifying the fibrotic phenotype (Figure 1C).

**Figure 1:**
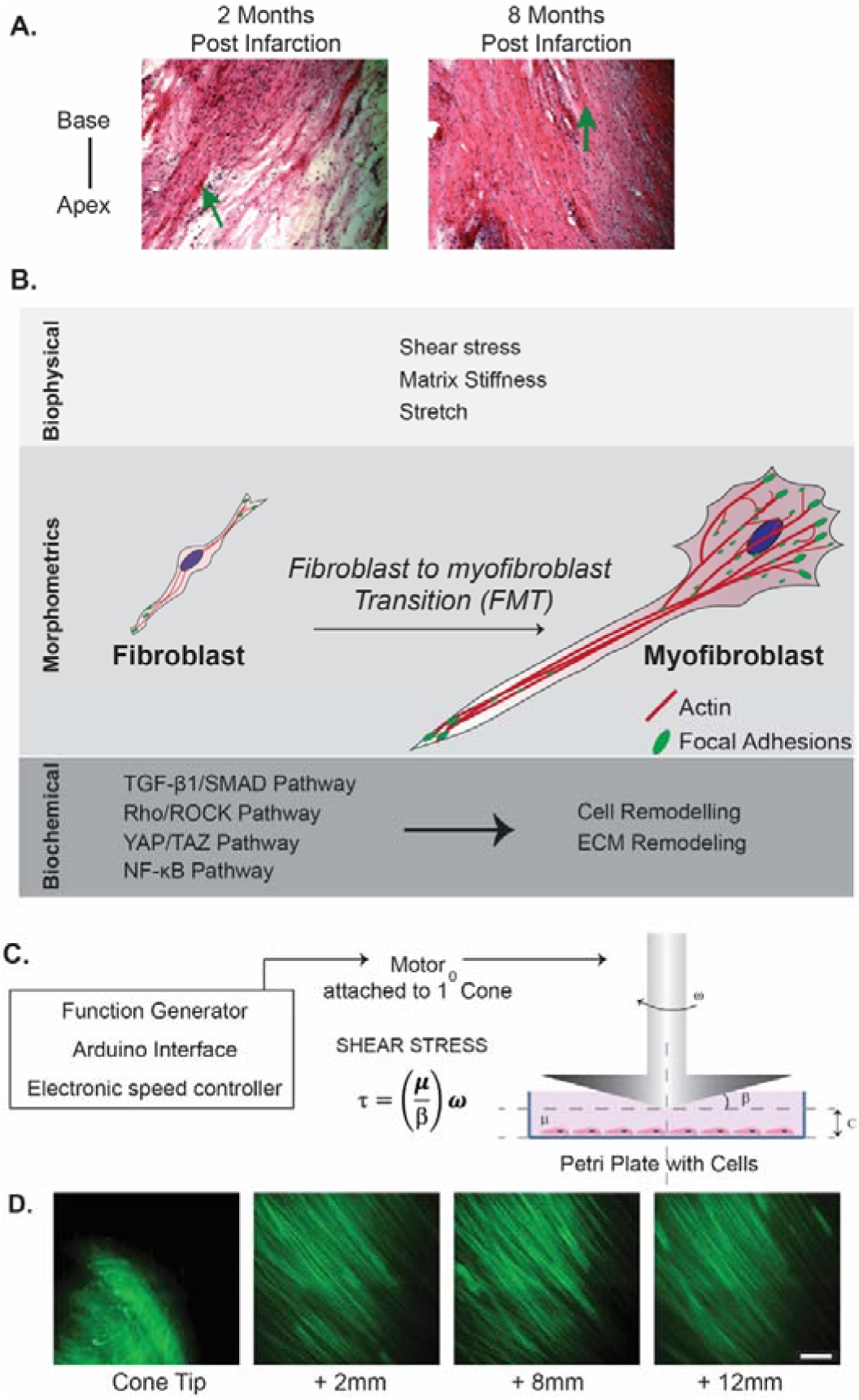
Stress-Induced Fibroblast to Myofibroblast Transition (FMT). (A) Representative histological sections of ovine myocardial tissue at 2 and 8 months post-infarction show significant changes in the mid-wall region^4^. Green arrows indicate remodeled fibrotic tissue characterized by a high density of fibroblasts. (B) Schematic shows the biophysical, morphological, and biochemical drivers underlying FMT. (C) A custom microscope-mounted fluid shear device was used to apply uniform FSS on fibroblasts over a prolonged duration. (D) Flow visualization studies demonstrate a large annual region of laminar shear that extends from 2 -12 mm from the 1° cone tip.

Mechanical forces are increasingly recognized as potent modulators of cellular behavior, influencing morphology, gene expression, and differentiation^10^. This recognition has led to a growing interest in understanding how mechanobiological cues contribute to the initiation and progression of fibrosis. Excessive dynamic stretching stimulates myofibroblast activation through the development of contractile actomyosin bundles, α-SMA expression, and stretch-dependent signaling^11^. Myofibroblasts have well-defined adhesions and higher expression of intracellular gap junctions^12^. Substrate stiffness also plays an important role in the trans-differentiation of fibroblasts into myofibroblasts, also called fibroblast-to-myofibroblast transition (FMT)^13,14^. Myofibroblast differentiation is mediated by TGF-β activity through its binding to receptors on the cell membrane; this activity is elevated on stiffer substrates and correlates with *α*5 integrin expression^13–15^. Recent studies show that cues from the three-dimensional collagen network structure regulate myofibroblast differentiation under static conditions^16^. Fluid shear stress (FSS) can also induce FMT: fibroblasts on 2D substrates elongate and align in the direction of shear, and differentiate into myofibroblasts under shear^17,18^. Low fluid shear (0.1-0.3 dyn/cm²) in 3D matrices induces alignment of the underlying collagen fibers and facilitates FMT through the upregulation of TGF-*β*1^19^. Fibroblast migration is enhanced through the interplay between shear stress and biochemical signaling pathways, including TGF-β, Rho/ROCK, FAK, and YAP, which promote cytoskeletal reorganization and drive the formation of contractile stress fibers^20,21,14,22,23^, yet the evolution of traction forces over time under continuous shear stress exposure remains incompletely understood. Despite advances in understanding FMT, the molecular mechanisms driving these processes due to physiological FSS remain poorly understood^24^. Significant gaps also remain in our understanding of how temporal sequences in mechanotransduction pathways driving FMT are initiated and orchestrated under prolonged fluid flows. Finally, there is little evidence to show if sheared cells retain a ‘mechanical memory’ of the transition once the shear stimulus is removed.

Our study aims to address these questions using a custom microscope-mounted fluid shear device (Figure 1D).^25–27^ The instrument enables precise control of the magnitude and duration of shear stress, permits real-time cell visualization, and provides a physiologically relevant milieu to identify mechanotransduction pathways during FMT. We applied laminar shear stress (1.2 Pa) to NIH3T3 fibroblasts and analyzed the temporal changes in traction stresses, migration, adhesion strength, and gene expressions during shear over 12 hours, and recovery post-shear for 12 hours. We demonstrate myofibroblast transitions induced due to prolonged physiologically relevant FSS, marked by increased gene expressions linked to focal adhesion, cytoskeletal, and ECM remodeling. Transformed cells exerted higher tractions and had enhanced adhesion strengths which persisted 12 hours post-shear. The interplay between shear stress and biochemical pathways promotes cytoskeletal reorganization and drives the formation of contractile stress fibers, enhancing fibroblast migrations. These results provide insights into fibrotic mechanisms, highlighting potential pathways for therapeutic intervention.

## RESULTS

### Time-dependent adaption of cytoskeletal and focal adhesions to shear stress

We hypothesized that prolonged FSS induces stress fiber and focal adhesion remodeling in fibroblasts. To study the effects of prolonged shear on cytoskeletal remodeling, we subjected NIH3T3 fibroblasts to constant shear stress of 1.2 Pa at 20 minutes, 1, 3, 6, and 12 hours using a custom microscope-mounted fluid shear device (Figure 1C; see Materials and Methods). The device features a 1*°* cone that rotates above a cell-seeded substrate using a hard drive motor, controlled with an electronic speed controller, generating a broad annular region of laminar shear stress (Figure 1D). This region of uniform FSS, extending 2 to 12 mm away from the cone tip, allows simultaneous assessment of a large number of cells in a single experiment over an extended duration. We stopped the shear stress after 12 hours (Shear group) and monitored cells over 12 hours in the absence of FSS (Recovery group). Cells were fixed at various time points and stained with rhodamine-phalloidin (red) to visualize actin, DAPI (blue) for the nucleus, and antibodies against zyxin (green) and vinculin (cyan) to identify mature and dynamic focal adhesions (Figure 2A). We used these images to assess time dependent changes in cell areas, aspect ratios, and stress fiber lengths due to prolonged FSS and post-shear, and during recovery (Figure 2B, C, D; see Materials and Methods).

**Figure 2:**
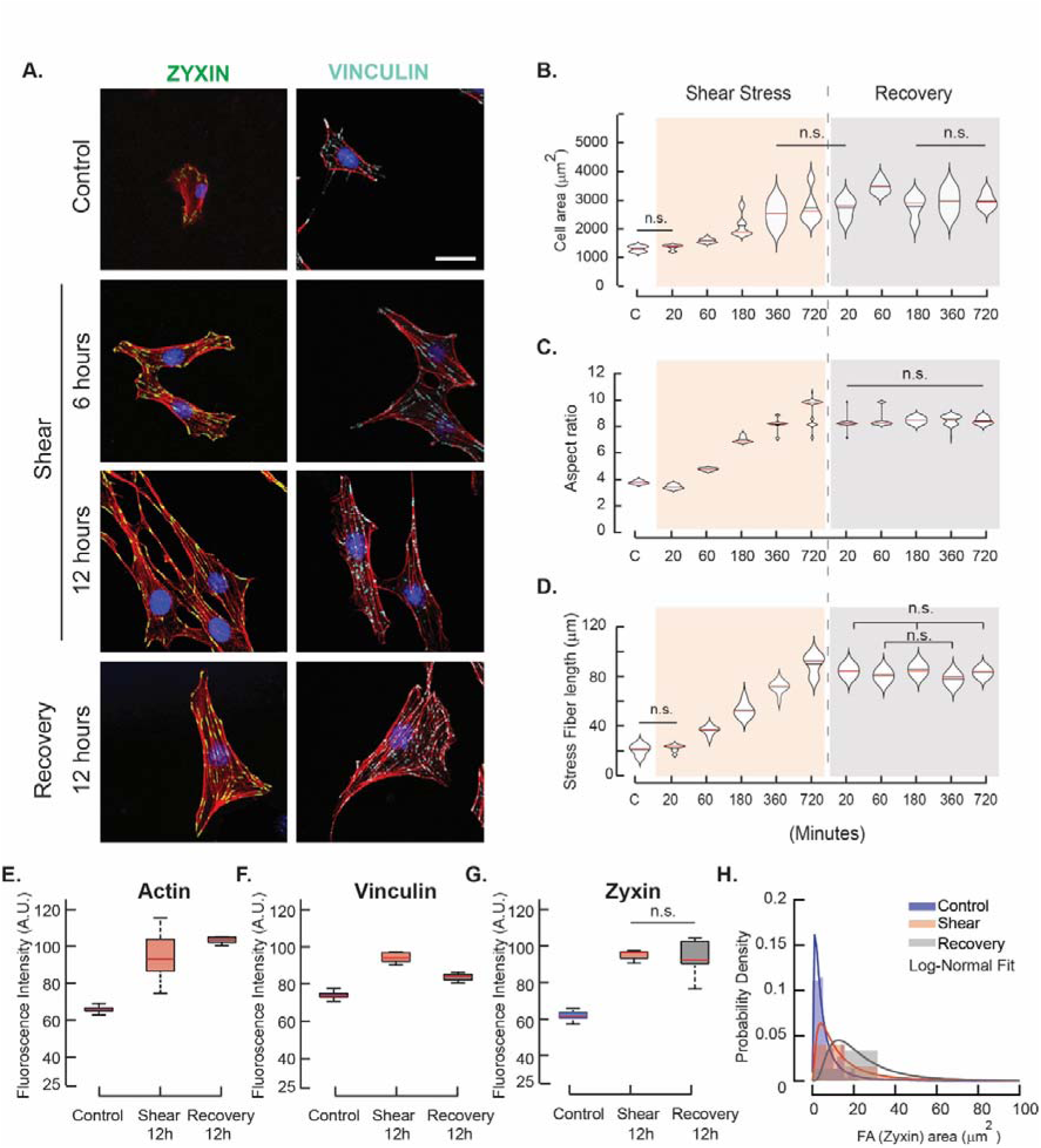
Time-Dependent Responses to Prolonged Shear Exposure. (A) Representative immunofluorescence images of NIH3T3 fibroblast cells from Control (no shear), 6 and 12 hours shear, and 12 hours post-shear (Recovery) groups. Cells were stained for actin (red), nucleus (blue), zyxin (green), and vinculin (cyan). Scale bar: 50µm (B) Variations in cell areas (n=50), (C) Aspect ratios (n=50), and (D) Stress fiber lengths were measured at each time point during shear and post-shear (n=15). Fluorescence intensities were quantified and compared for (E) Actin, (F) Zyxin, and (G) Vinculin in the focal adhesions for Control, (12 hours) Shear, and (12 hours post-shear) Recovery groups (n=15 each group). Differences between the non-significant groups alone are labeled; all other group comparisons show statistically significant differences at p<0.05. (H) Probability density plots with log-normal fits are shown for projected zyxin areas (µm²) from the images.

Our data demonstrate distinct time-dependent changes in cell morphologies during FSS application (Figure 2A). Following shear, cells show prominent stress fiber with zyxin and vinculin localization. Cell areas gradually increased under FSS (Figure 2B), with no significant differences at 20 minutes compared to Control (no shear). However, the aspect ratios initially decreased before increasing steadily over time (Figure 2C). These results suggest that a minimum stress duration is required to induce observable changes (Figure S1). By 1 hour, cell areas increased significantly, indicating fibroblast remodeling, alignment, and elongation in the shear direction (p<0.05). Aspect ratios and cell areas plateaued between 6 and 12 hours of FSS (p<0.05) and remained unchanged in the Recovery group. The stress fiber lengths also increased in a time-dependent manner, peaking to 89.12L±L8.04Lμm at 12 hours before marginally decreasing during recovery (Figure 2D; see Materials and Methods). These findings suggest that prolonged FSS exposure induces cytoskeletal modifications that drive morphological changes.

We quantified changes in actin, zyxin, and vinculin intensities from the immunofluorescence images of cells (n=15) in the Control, Shear (12 hours), and Recovery (12 hours post shear) groups, respectively (Figure 2E-G; see Materials and Methods). Actin intensity was significantly higher in the Shear group compared to Control (p<0.05), indicating extensive cytoskeletal remodeling under FSS. These changes persisted 12 hours after the removal of the fluid stimulus. Vinculin and zyxin intensities, markers of mature and dynamic focal adhesions, also increased with shear duration, suggesting progressive strengthening of mature focal adhesions under fluid shear. The recovery group showed reduced vinculin intensity, compared to zyxin-positive regions that were unchanged after the removal of FSS (Figure 2F, G). The distribution of focal adhesion areas, fit to a log-normal model, showed smaller adhesions in the Control group compared to the Shear and Recovery groups that had larger focal adhesions (Figure 2H). The observed focal adhesion growth and increased stress fiber lengths under prolonged FSS indicate significant changes in biophysical properties that remain elevated for 12 hours post-shear.

### Prolonged shear induces FMT, and enhances cell contractility and adhesion strength

How does stress fiber and focal adhesion remodeling alter the biophysical properties of cells? To investigate this, we seeded fibroblasts on bead-coated polyacrylamide substrates (10 kPa) and applied 1.2 Pa shear using the custom microscope-mounted device (see Materials and Methods). Figure 3A shows representative brightfield images alongside corresponding traction maps, calculated using a regularized Fourier Transform Traction Cytometry (Reg-FTTC) method^28^, from the Control, Shear (12 hours), and Recovery (12 hours) groups, respectively. These data provide the first traction measurements of cells subjected to prolonged elevated FSS.

**Figure 3:**
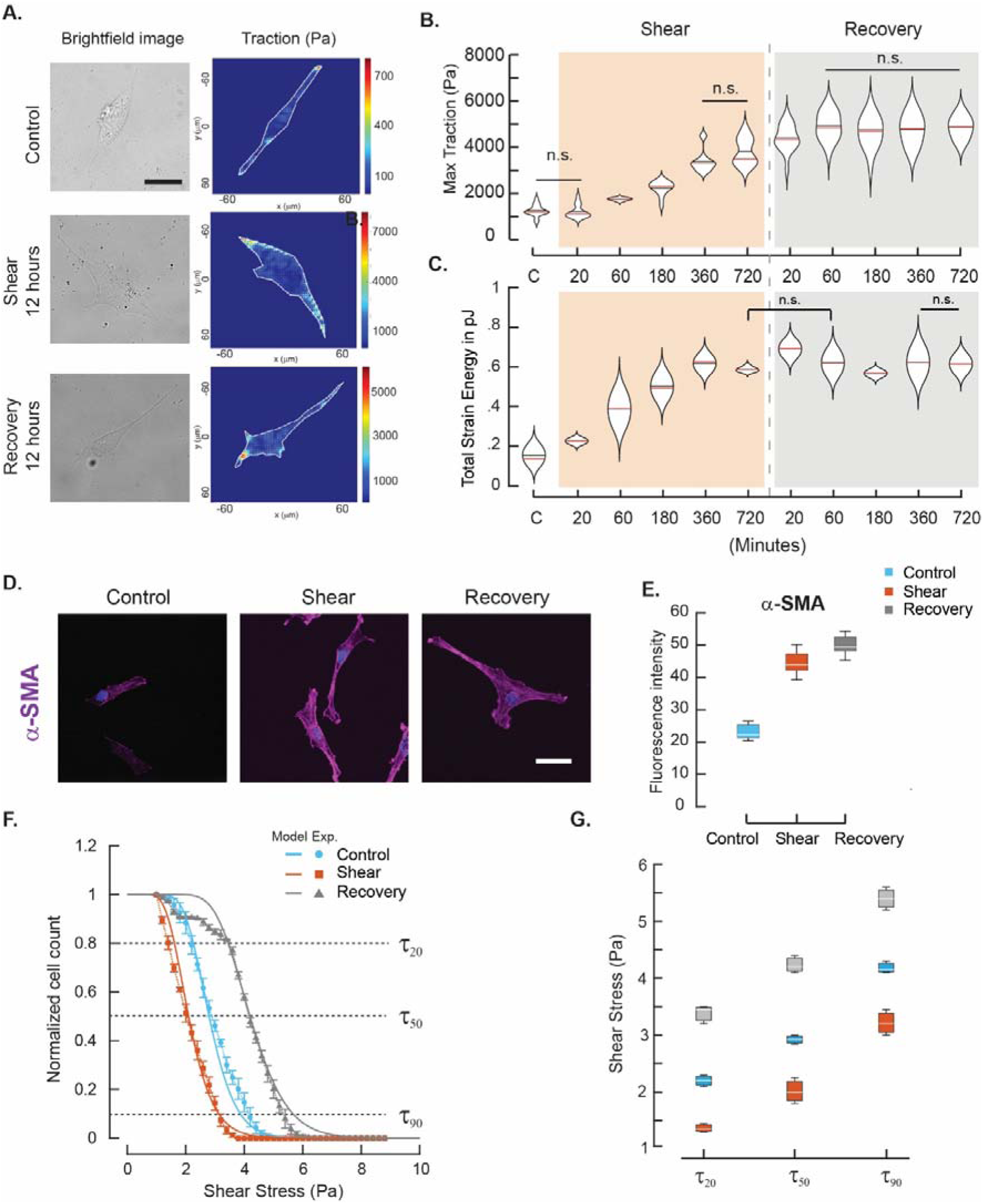
Cell Contractility and Adhesion Dynamics under FSS and post-shear. (A) Representative brightfield images and corresponding traction stresses (Pa) are shown from Control, Shear, and Recovery groups measured using the regularized Fourier Transform Traction Cytometry (Reg-FTTC) method^28^. Scale bar: 50µm. (B) Maximum tractions increased from 60 to 360 minutes during shear and did not change after removing the shear (n=15). (C) Strain energy increased with time due to shear and remained relatively unchanged post-shear (n=15). (D) Immunofluorescence images of α-SMA (purple) demonstrate activation of mechanotransduction pathways. Scale bar 50µm. (E) Quantification of α-SMA expression across Control, Shear, and Recovery conditions with fluorescence intensity (A.U.). (F) The de-adhesion strengths of cells are reported for cells in Control (n=370), Shear (n=366), and Recovery (n=375) groups. Normalized cell number followed sigmoidal behaviors (G) Clear differences are visible between the weakly adherent (*τ*_20_) shear stresses, critical deadhesion strengths (*τ*_50_), and strongly adherent (*τ*_90_) shear stresses. Non-significant differences between the groups are indicated using n.s. All other groups are statistically significant (p<0.05).

Maximum tractions and total strain energy densities increased steadily under shear (Figure 3B, 3C), with higher tractions localized at the polarized cell ends. Tractions increased significantly after ∼ 1 hour of shear (p<0.05) but showed no further differences between the 6 hours and 12 hours Shear groups (p<0.05). Surprisingly, tractions increased significantly and remained unchanged after the removal of the shear stimulus, indicating that increased contractile activity persisted during recovery (p<0.05). Corresponding strain energies also increased with prolonged FSS. Figure 3C shows a marginal increase from 20 minutes to 1 hour, followed by a pronounced increase between 3 and 12 hours. However, these values remained elevated post-shear in the Recovery group. Stable focal adhesions (Figure 2A) and stress fibers (Figure 2D) support a highly contractile state, which is associated with increased α-SMA expression — an indicator of the contractile phenotype in fibroblasts. α-SMA was significantly elevated under FSS and remained unchanged after shear removal (p<0.05; Figures 3D, E). Sustained tractions and cytoskeletal changes suggest retention of “mechanical memory” and a potential shift towards a myofibroblast-like phenotype, which may influence cellular remodeling and fibrosis progression.

We hypothesized that prolonged shear alters the adhesion strengths of transformed cells. To test this, we measured the critical adhesion strengths of cells in the Control, Shear, and Recovery groups using the device on fibronectin-coated substrates (see Materials and Methods). The FSS was steadily ramped from 1.2 Pa in increments of 0.2 Pa/ minute in these experiments until all cells detached from the substrate (see Materials and Methods section)^24,26^. The resulting sigmoidal graph (Figure 3F) shows the fraction of cells remaining on the substrate with each step shear increase. We quantified the critical deadhesion strengths (*τ*_50_) using these data and obtained the stresses required to detach weakly adherent (*τ*_20_) and strongly adherent cells (*τ*_90_) from the substrate. The rightward shift in deadhesion curves post-shear indicates increased adhesion strength. Interestingly, significant deviations in the initial part of the deadhesion curve for the Recovery group suggest that ∼15% of cells retained highly dynamic adhesions, similar to the Control group. In contrast, the Shear group exhibited a leftward shift relative to Control, suggesting weaker and more dynamic adhesions (Figure 3G).

We fit the deadhesion curves to a cell detachment model incorporating experimentally obtained fibroblast area distributions (Equation 1) ^25,27^. The model simulates stochastic binding kinetics of integrin-mediated slip bonds, distributed around the periphery of hemispherical cells. Figure 3G shows that the model effectively captures these experimental results. However, the Recovery group exhibited significant deviations from the model, particularly at *τ*_10_ and *τ*_90_ indicating high adhesive heterogeneity in the cell population post-recovery. This heterogeneity likely arises from variations in myofibroblast transitions and potential reversals^29^. These deadhesion dynamics highlight the complex interplay between adhesion strength and cytoskeletal adaptation to prolonged shear stress.

### FSS induces changes in gene expressions for cytoskeletal, focal adhesions, and ECM proteins

To explore the signaling pathways driving fibroblast adaptation to prolonged FSS, we analyzed mRNA levels of key genes in extracellular matrix (ECM) remodeling, focal adhesion dynamics, and the cytoskeletal regulation across Control, Shear, and Recovery groups. We selected primers targeting genes associated with actin dynamics (Actin, RhoA), ECM (Collagen-1, Collagen-3, Fibronectin, MMP2, MMP9), focal adhesions (Vinculin, Talin, FAK, IGTAV, IGTA5, IGTB1, IGTB3), and α-SMA, a myofibroblast differentiation marker (see Materials and Methods). Gene expressions were normalized to GAPDH levels and the Control group.

Figure 4A shows distinct time-dependent changes in gene expression profiles corresponding to the ECM, triggered by prolonged FSS. Fibronectin expression significantly increased, peaking between 3 and 12 hours of FSS, before reducing post-shear. Collagen-1 and Collagen-3 — key structural components of the ECM — were upregulated during shear and in the recovery phase. Matrix metalloproteinases corresponding to MMP2 and MMP9 displayed distinct changes under shear: MMP2 levels remained elevated from 3 hours of FSS through the post-shear phase, whereas MMP9 levels were higher post-shear. Focal adhesion-related genes for Vinculin and Talin were minimally upregulated at 3 and 6 hours of shear (Figure 4B), aligning with observations on maximal cell elongation and flow-induced alignment. Integrins IGTAV and IGTA5 showed a moderate increase during shear exposure but were significantly upregulated during recovery. IGTB3 exhibited a steady and gradual increase through the shear exposure, suggesting its involvement in stabilizing adhesions under prolonged FSS.

**Figure 4:**
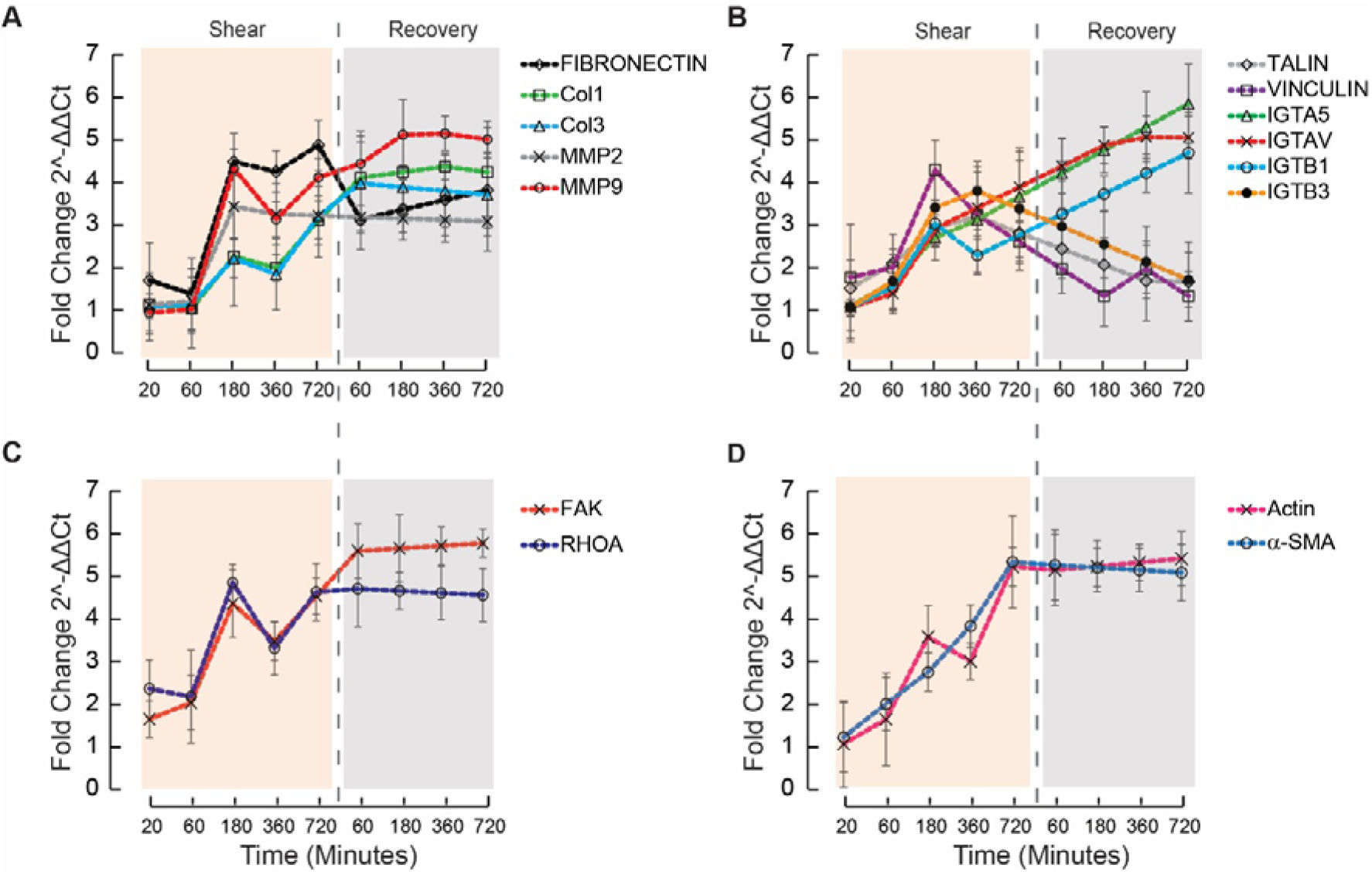
Gene Expression Changes under Prolonged Shear. Fold changes in gene expression levels (2^-ΔΔCt^) are shown during a steady increase in FSS and recovery phases. Primers corresponding to (A) key extracellular matrix (ECM) constituents (Fibronectin, Collagen I (Col1), Collagen III (Col3), MMP2, and MMP9). (B) Focal adhesion markers including Talin, Vinculin, ITGA5, ITGAV, ITGB1, and ITGB3. (C) Signaling molecules (FAK and RhoA) and (D) myofibroblast markers (Actin and α-SMA) show clear changes.

Shear stress significantly upregulated RhoA and FAK expressions by 3 hours (Figure 4C), with elevated levels persisting over 12 hours post-shear. Increased RhoA expression suggests active cytoskeletal reorganization under shear, consistent with the observed increases in traction stress and stress fiber formation. Similarly, FAK, a central mechanotransduction regulator, was elevated alongside actin and α-SMA, both of which increased steadily under shear. These data indicate cytoskeletal strengthening and myofibroblast transition which persisted during recovery, indicating the transformation of fibroblasts into myofibroblasts that have stable adhesions and prioritize ECM deposition. Our results highlight the complex regulatory mechanisms governing fibroblast adaptation to shear stress, emphasizing immediate and sustained signaling changes that drive fibroblast transformation under prolonged fluid shear.

### Migration and directionality of fibroblasts

Fibroblast migration is essential in early fibrosis for tissue repair. In contrast, myofibroblasts are characterized by increased contractility, and ECM deposition to remodel tissues. We hypothesized that prolonged FSS enhances directed fibroblast migration but reduces myofibroblast motility. Figure 5A-C shows rose plots of migration behaviors in Control, Shear (12 hours), and Recovery (12 hours) conditions. Under FSS, fibroblasts migrated along their oriented directions (Movies 1-3). In contrast, cells in the Recovery group surprisingly exhibited minimal directional changes, migrating along straight paths (Figure 5C). Mean squared displacement (MSD) was significantly higher in the Shear group (Figure 5D), driven by cytoskeletal remodeling and dynamic focal adhesion formation under FSS (see Materials and Methods). In contrast, MSD values were significantly lower in the Recovery group, suggesting stable adhesions, as myofibroblasts prioritize ECM deposition over migration. Thus, FSS enhances fibroblast mechanoadaptability, while myofibroblasts reinforce adhesion stabilization, shifting to a less migratory phenotype. The effective diffusivity (α), derived from the slope of the log(MSD) with log(time) data (Figure S3), was 0.81 in the Control group, indicating sub-diffusive migration without directional cues. In contrast, α increased to 1.26 in the Shear group, reflecting super-diffusive migration with persistent flow-aligned movement. The Recovery group had a lower α (=0.59), signifying constrained, sub-diffusive migration with loss of directional ability.

**Figure 5:**
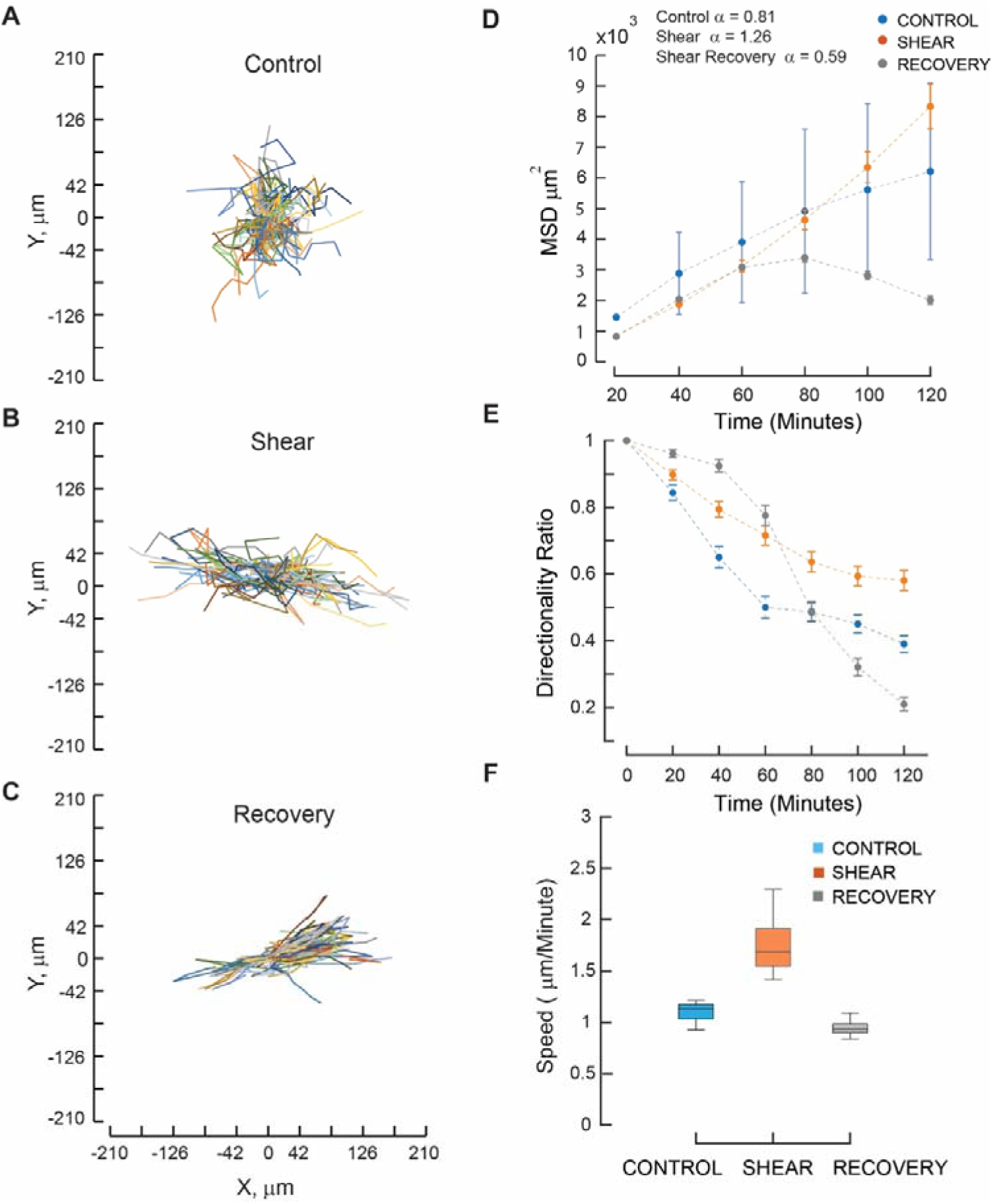
Cell Migration Dynamics under FSS and during Recovery. Migration tracks of cells from (A) Control (n=87), (B) Shear (n=90), and (C) Recovery after removal of shear stimulus (n=98) illustrate displacements over 2 hours that have different dynamics. (D) Mean squared displacement (MSD) highlights differences in migrations between groups. (E) Directionality ratios indicate changes in the orientation of cell migration. (F) Cell migration speeds (µm/min) were significantly different in comparisons between the Control, Shear, and Recovery groups (p<0.05).

Directionality Ratios (DR) were significantly higher in the Shear group than in the Recovery group at 2 hours, indicating sustained directional persistence along specific directions post-shear (p<0.05; Figure 5E). However, myofibroblasts had significantly lower DR values within ∼60 minutes post-shear. Migration speeds also differed significantly between all groups (p<0.05; Figure 5F). These differences highlight the role of FSS on cytoskeletal dynamics and adhesion strengths in fibroblasts, whereas myofibroblasts retained their mechanical memory of the transformation. Prolonged shear stress hence induces fibroblast-to-myofibroblast transformation, leading to stabilized adhesions, reduced motility, and elevated contractility (Figure 3B). These results hence show fibroblast mechano-adaptivity as a critical factor in tissue repair, remodeling, and the progression of fibrosis.

## DISCUSSION

Fibroblasts are of mesenchymal origin and regulate the balance between ECM synthesis and breakdown to maintain tissue homeostasis^30^. Disruption of this fine balance in fibrotic disease stimulates the differentiation of fibroblasts into myofibroblasts due to chemokine signaling and mechanical activation^24^. While initially protective, FMT leads to excessive ECM deposition, tissue stiffening, and impaired organ function in fibrotic disease^31,32^. We investigated fibroblast adaptation to prolonged physiological shear stress (1.2 Pa), and its subsequent recovery post-shear, and uncovered key mechanisms in the complex and nonlinear coupling between biochemical signaling and biophysical changes driving FMT. Our study highlights four key findings: first, fibroblasts adopt an elongated spindle-shaped myofibroblast morphology under shear that is linked to increased stress fiber lengths and altered distributions of vinculin and zyxin along the cell periphery. These changes remained unchanged 12 hours post-shear, emphasizing the presence of cellular mechanical memory. Second, FSS induced cytoskeletal remodeling increases cell tractions in myofibroblasts, and upregulates actin and α-SMA expressions, sustaining the contractile phenotype post-shear. Third, prolonged FSS reduced the critical deadhesion strengths, and induced significant, time-dependent changes in ECM-related genes (FIBRONECTIN, Col1, Col3, MMP2, and MMP9) and cytoskeletal remodeling (Actin, *α*-SMA, RHO), indicative of the myofibroblast transition^33,34^. Upregulation in RhoA, FAK, actin, and α-SMA expressions due to shear highlight FMT-associated mechanotransduction in fibroblasts. Finally, prolonged shear enhances fibroblast migrations (MSD, directionality, and speed), demonstrating distinct mechanical adaptations that are crucial for effective tissue repair in fibrosis.

The disruption of “function dictates form” provides a powerful framework for understanding how tissues adapt to forces that drive disease progression. Polarized, spindle-shaped fibroblasts aligned with flow due to increased stress fiber lengths^35^. This cytoskeletal reorganization, mediated by actin and α-SMA, is accompanied by changes in vinculin and zyxin expression. Shear-induced upregulation of α-SMA, a hallmark of FMT, begins ∼6 hours of shear exposure and persists post-shear. The accompanying focal adhesion growth, reorganization, and reinforcement further support the transition^36,37^. Myofibroblasts have microfilament bundles that terminate at specialized mature adhesion complexes, facilitating ECM attachment and contractility^24,30,38^. Stress fibers contribute to the formation of mature adhesions, enabling cells to resist FSS by generating tractions^10,19,32^.

Focal adhesion reinforcement and stress fiber formation are both critical in maintaining the contractile myofibroblast phenotype^39,40^. Studies have shown that traction forces play a crucial role in FMT, demonstrating that increased contractility, focal adhesion remodeling, and mechanotransduction pathways drive myofibroblast differentiation in fibrosis and wound healing; however, how prolonged shear stress sustains this transition has not been shown^41–44^. Despite marginal decreases in stress fiber lengths and vinculin expression post-shear, myofibroblasts retained the ability to generate elevated tractions. Increased zyxin in fibroblasts (FigureL2G, 2H) further supports its role in strengthening and stabilizing the cellular architecture^37,45,46^.

Shear induced upregulation of RhoA and FAK regulates cytoskeletal dynamics through biochemical signaling, enhancing cellular contractility and altering the tensional homeostasis^34^ ^47^. Elevated expression of ECM-related genes, such as fibronectin, collagen-1/3, and matrix metalloproteinases (MMP2 and MMP9), in the Shear and Recovery groups indicates active tissue remodeling, a hallmark of FMT^18^. MMP2 regulates ECM degradation and remodeling^48^, whereas MMP9 helps breakdown the ECM under stretch^49^. Notably, MMP2 expression remained elevated through shear exposure and recovery, whereas MMP9 showed a biphasic response, peaking at 3 and 12 hours. These findings align with gene expression changes observed during tissue remodeling in myocardial infarction, lung fibrosis, and wound healing^30^. Focal adhesion remodeling and stress fiber reinforcement during shear and post-shear altered the migration dynamics of fibroblasts. Directionality ratios and migration speeds significantly increased in the Shear group, which exhibited sub-diffusive migration compared to super-diffusive behaviors of Control cells (Figure 5D-F). These results agree with reports that show fibroblasts lack directionality under static conditions and have persistent anomalous migration^17^. Prolonged shear induced cellular polarization, with cells predominantly migrating along the shear direction in our study.

There are some limitations of this study: first, collagen fibers align under shear, creating tracks that facilitate cell elongation and migrations along the fiber direction. Figure S4 shows fluorescence images of collagen-functionalized coverslips subjected to uniform shear for 30 minutes in the device, demonstrating fiber alignment that may aid the realignment responses of fibroblasts. However, isolating the individual contributions of shear and collagen alignment remains challenging. Second, the observed mechanical memory of cells at 12 hours post-shear may not persist over longer durations, warranting further investigation. Third, the media used to cause fluid shear on cells may contain released soluble factors that contribute to FMT. We did not however assay the media for the presence of such chemicals. Finally, we did not investigate potential changes in proliferation and apoptotic markers under shear. Understanding the mechanotransductive processes driving FMT is crucial for developing therapeutic strategies to mitigate fibrotic diseases, where excessive myofibroblast activity leads to pathological tissue remodeling and stiffening. Targeting myofibroblast mechanoadaptation offers a promising avenue for the management and treatment of fibrosis.

### SIGNIFICANCE STATEMENT

Fibroblasts play a critical role in the synthesis, organization and extracellular matrix remodeling, contributing to tissues homeostasis. Their trans-differentiation into myofibroblasts influences tissue remodeling and fibrotic progression. How does prolonged shear impact fibroblast mechanobiology? We subjected fibroblasts to laminar shear using a custom device and measured temporal changes in cell signaling and biophysical properties. We demonstrate that sustained shear drives fibroblast-to-myofibroblast transition through progressive actin and α-SMA remodeling, leading to the formation of elongated stress fibers, and increased vinculin and zyxin expression which remains elevated post-shear. Myofibroblasts have increased traction stresses that persisted post-shear, suggesting a mechanical memory of the transformation. Deadhesion strengths increased significantly post-shear, accompanied by shifts in gene expressions governing extracellular matrix homeostasis, which affect migration dynamics.

## ACKNOWLEDGEMENTS

We acknowledge the Bioimaging Facility, Indian Institute of Science (IISc) for help with the use of the Andor Dragonfly 502 spinning disk confocal microscope. NG is grateful to DBT (BT/BR23724/BRB/10/1606/2017) and SERB (SPG/2021/002425) for project support. The CFX96 Touch Real-Time PCR Detection System in the Biomechanics lab was acquired through a generous grant from DST-FIST. We also thank Prof. Pramod Pullarkat (RRI, India) and Prof. Gautam Menon (Ashoka University, India) for useful discussions during the early stages of this work.

## Author contributions

NP: Methodology, Software, Validation, Formal Analysis, Investigation, Writing – Original Draft, Data Visualization. AJ: Methodology, Software, and Validation of migration result. PK: Conceptualization, Methodology, Formal Analysis, Writing – Reviewing and Editing, Supervision. NG: Conceptualization, Methodology, Software, Formal Analysis, Resources, Data Curation, Writing – Original draft, Reviewing and Editing, Data Visualization, Supervision, Project Administration and Funding Acquisition.

---

**Supplementary Figure S1:**
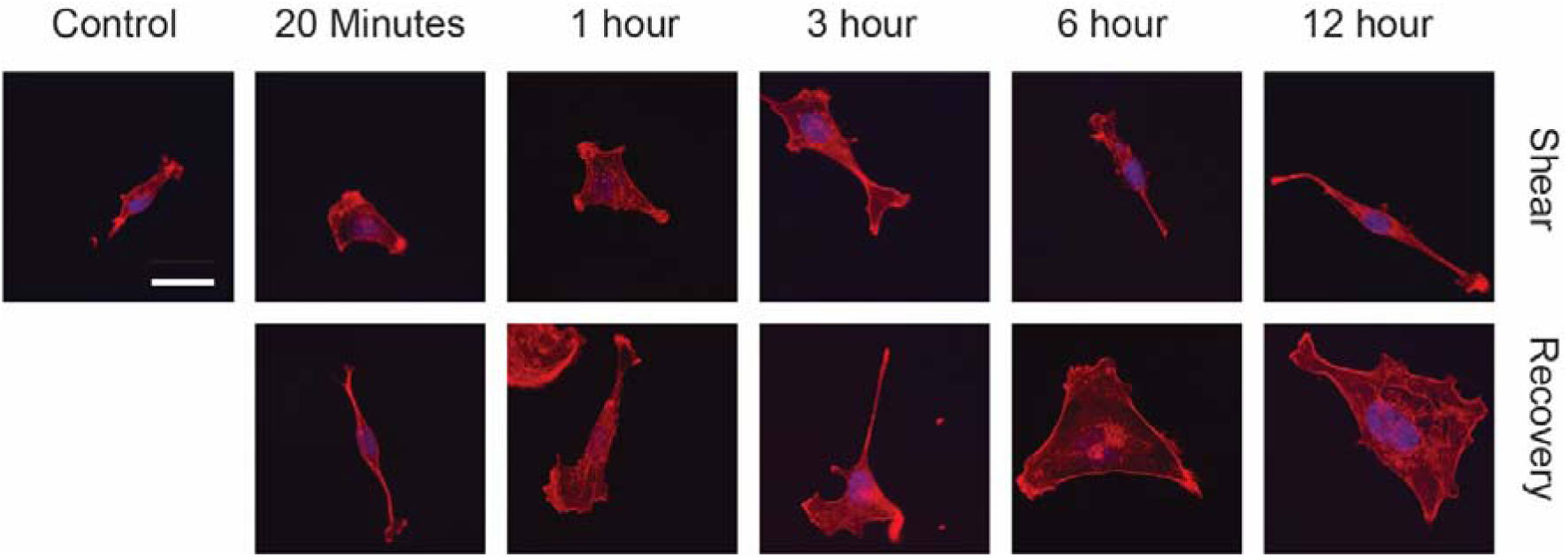
Representative actin and DAPI stained images of fibroblasts at various points during shear and recovery. Shear direction from bottom to top. Scale 20um.

**Supplementary Figure S2:**
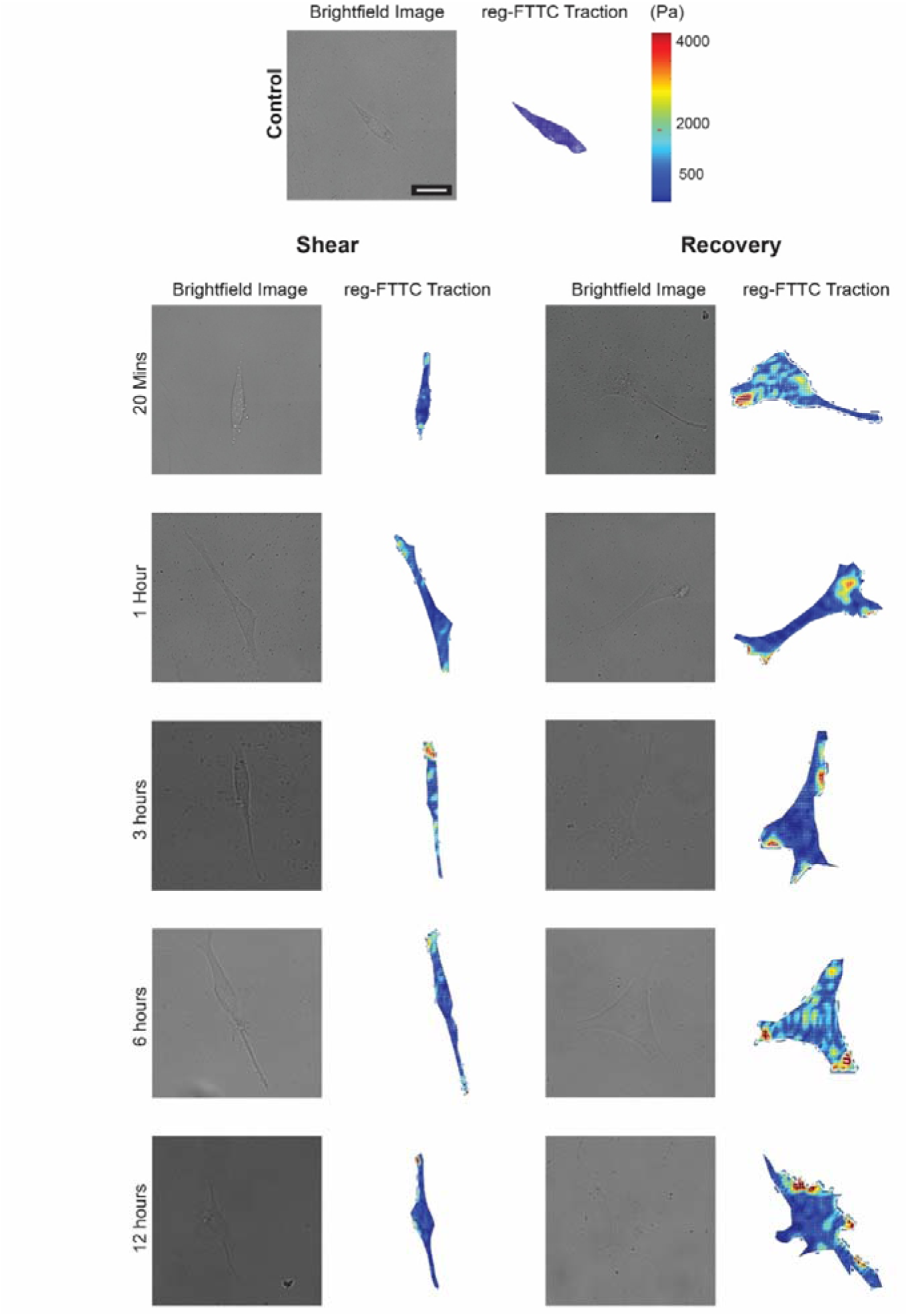
Brightfield and traction maps of representative cells at various points during shear and recovery. Tractions, calculated using a regularized FTTC method, show higher values at the polarized ends of the cell for cells under shear. Scale 20 *μ*m.

**Supplementary Figure S3:**
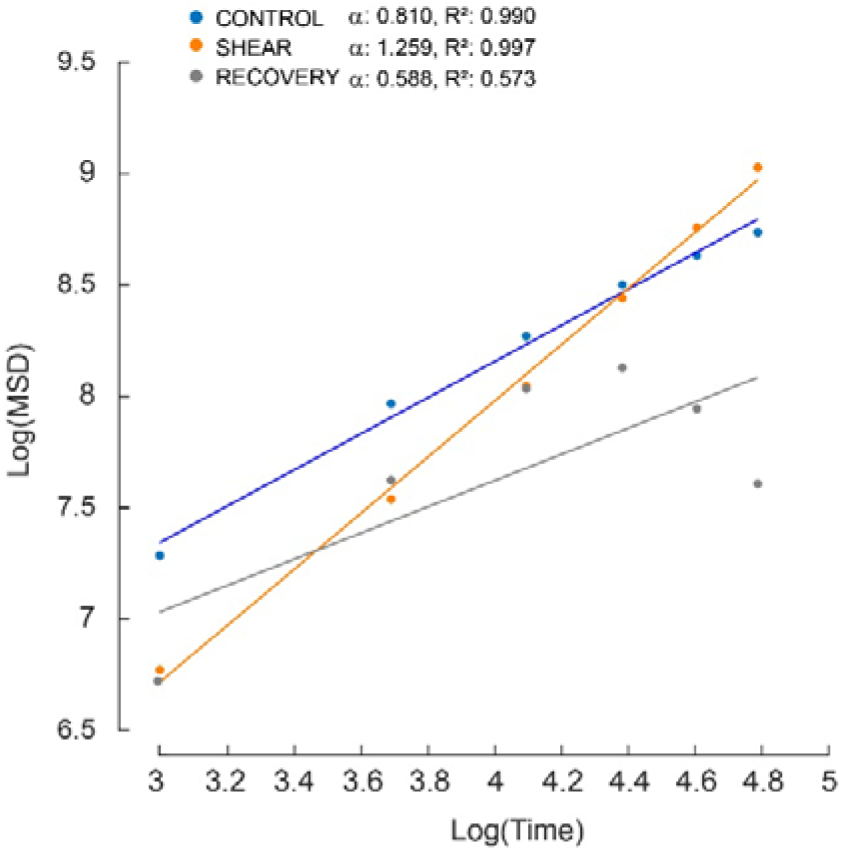
Mean Squared Displacement (MSD) variations with time are shown in log-log plots for cells in Control (blue), Shear (orange), and Recovery (gray) groups. The slopes to these data, *α*, are the effective diffusivity values that were used to compare with various diffusive behaviors. R^2^ values show the goodness of fit to the data.

**Supplementary Figure S4:**
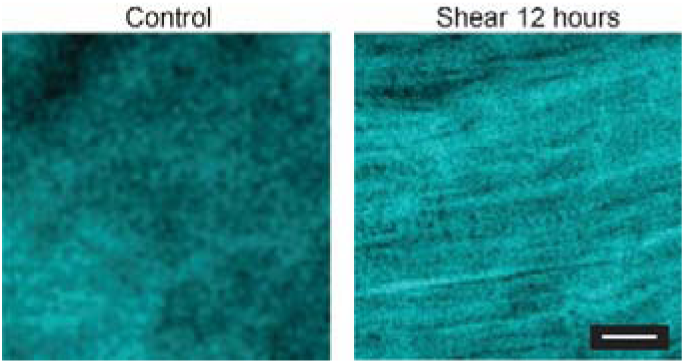
Collagen alignment on substrates under 1.2 Pa shear was determined using 40 *μ*g/ mL collagen I petridishes. Images are shown 12 hours post-shear. Scale bar: 20 *μ*m.

## Supplementary videos

**Movie 1:**
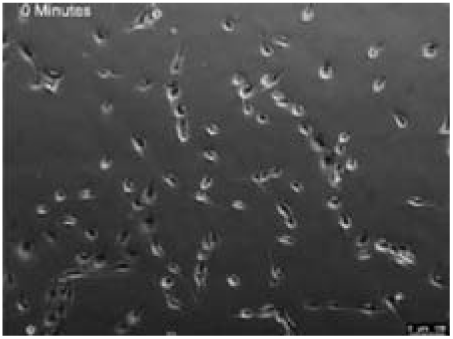
Control cell migrations on fibronectin coated substrates show persistent random migrations Time-lapse video shows cell migrations in the Control group over 2 hours. Cells exhibit active random migrations movement over time.

**Movie 2:**
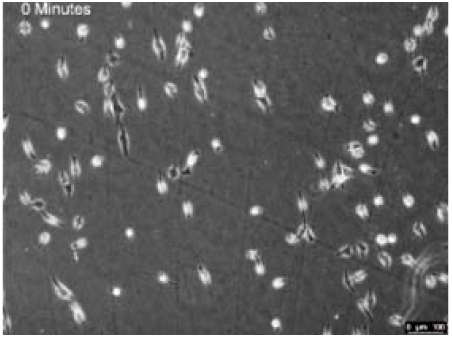
Sheared cells have directed migration on fibronectin-coated glass substrates Time-lapse video shows fibroblast migrations over 2 hours following a 12 hours exposure to 1.2 Pa FSS. Cells have increased alignment and directional persistence along the shear direction.

**Movie 3:**
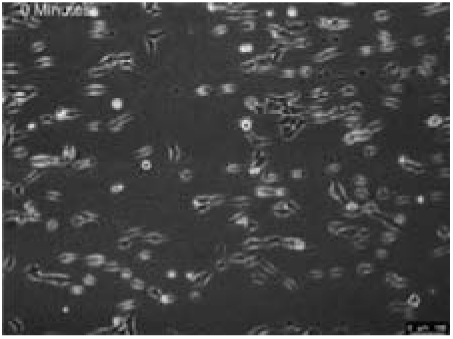
Removal of FSS showed migrations along straight trajectories over 2 hour period Transformed fibroblasts following 12 hours of shear exposure were maintained for 12 hours without shear, and migrations were quantified over 2 hours. Cells display reduced migration speeds and move primarily along one direction.

## MATERIALS AND METHODS

### Cell Culture

NIH-3T3 mouse embryonic fibroblast cells at early passage numbers were maintained in Dulbecco’s Modified Eagle Medium (DMEM, Invitrogen) supplemented with 10% (v/v) fetal bovine serum (FBS, Invitrogen) and 1% (v/v) penicillin/streptomycin (Sigma-Aldrich) in a humidified incubator at 37°C with 5% CO_2_. Cells were passaged every 2–3 days and harvested by washing thrice with phosphate-buffered saline (PBS) and detached from the flasks using 0.25% trypsin-EDTA (Invitrogen) for experiments. Fibronectin (40 µg/ml) was added to a coverslip at 37 °C for 1 hour in a humid chamber and washed thrice with PBS for experiments. Cells were seeded at 5000 cells/ml on fibronectin coated coverslips and allowed to attach and spread for 6 hours before starting the shear experiments.

### Fluid shear studies using a custom microscope-mounted device

A custom fluid shear device was used for all studies (Figure 1D). The device was placed on a live-cell epifluorescence microscope (Leica DMI 6000) equipped with a PECON cage incubator for live-cell imaging during shear. Details of the device are provided in our previous work^24,26^. Briefly, a 1° cone was rotated using a hard drive motor powered by a DC supply and controlled *via* an Arduino-based electronic speed controller to modulate pulse width and frequency (Figure 1D). Shear stresses, calculated using equations for Couette flows generated in the device, are directly proportional to the angular speed of the motor. The constant rotation of the cone produces a large annulus of laminar shear stress (Figure 1D). Flow visualization experiments, performed using fluorescently labeled beads (2 *μ*m size; 2 *μ*gm/ml), were used to identify the regions of laminar flow in the device. An annular laminar flow region of 2-12 mm, located away from the cone tip and ends of the Petri plate, was used in subsequent analyses.

Cells were stained with Hoechst (Invitrogen) for 3 minutes in an incubator maintained at 37 °C and 5% CO₂ and plated on fibronectin coated coverslips in the device. Uniform 1.2LPa FSS was exerted on the cells and the experiments were stopped at various time points during the application of shear for cell morphometry studies and to quantify changes in the gene expressions. These intervals included 20 minutes, 1, 3, 6, and 12 hours. We also quantified changes in the critical deadhesion strength of fibroblasts based on earlier protocols^24,26^ by ramping the shear stress by 0.2LPa per minute using a custom Arduino code. Images acquired using a CCD camera at each stress increment were used to count the number of cells on the substrate based on the fluorescence of the Hoechst-stained nuclei. The ratio of cells remaining (N/N_₀_) on the substrate showed sigmoidal dependence with shear stress. The critical adhesion strength (τ_₅₀_) was defined as the shear stress at which 50% of the cells remained adhered to the substrate. Shear stresses based on a 20% cell fraction removed from the substrate, *τ*_20_, was calculated to estimate weakly adherent cells. *τ*_₉₀_ value of shear stress was used to characterize strongly attached cells based on the removal of 90% of cells from the substrate^24^.

Experimental sigmoidal curves were fit to a cell detachment model based on the statistical distribution of cell areas and adhesion strengths^24^. The cell was approximated as a hemispherical shape which was attached to a substrate using a uniform distribution of focal adhesions along the cell perimeter. This model considers heterogeneity in cell size and adhesion strength to calculate the fraction of cells adherent on the substrate at each step increase in shear stress due to Stoke’s flow in a low Reynold’s number moving fluid. Bell’s deterministic equations were used to model the molecular clusters as slip bonds which are stable with an infinite lifetime until they experience a critical load. The fraction of cells remaining on the substrate, Φc.J, is given as:

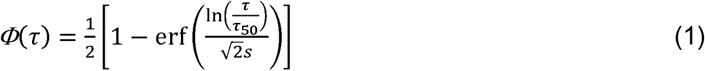

In this expression, the parameter , is proportional to the distribution width of attachment points for cells, as determined from experimental data of cell spread areas. The term ._50_ represents the critical shear stress required to detach 50% of the cells from the substrate, and *Φ*(.) denotes the fraction of adherent cells at a given shear stress .. Data from experiments were fit to the model using MATLAB (v R2019b, The Mathworks Inc. Natick, MA).

### Traction Force Microscopy (TFM)

We assessed changes in cell contractility under shear stress using traction force microscopy (TFM), described in our earlier work^28^. Briefly, glass coverslips were cleaned and prepared with polyacrylamide (PA) gels of 10 kPa stiffness using a mixture of acrylamide and bisacrylamide in distilled water, combined with 10% APS and TEMED. An optimized fluorescent bead density on top of the gel surface allowed the use of an epifluorescence microscope for the traction studies. The PA gels were functionalized with sulpho-SANPAH and coated with fibronectin (40 *μ*g/ml) for cell attachment. NIH3T3 cells were seeded onto the gels at 2000 cells/ml and allowed to attach for 6 hours in an incubator at 37 °C with 5% CO_₂_. The substrate with cells on PA hydrogel was placed in the custom device and subjected to 1.2LPa shear stress for various durations (20 minutes, 1, 3, 6, and 12 hours) and during recovery (n=15 in each group). Phase-contrast/ epifluorescent images of the cells and beads were acquired using a Leica DMI6000B live-cell microscope equipped with a PECON stage to obtain the cell boundary and deformed images of beads on the hydrogel surface due to cell attachments. Cells were removed using trypsin-EDTA, and the referential bead positions were acquired. Traction data were analyzed using a custom regularized Fourier Transform Traction Cytometry (Reg-FTTC) MATLAB code^28^.

### Migration studies

NIH3T3 cells from Control (no shear), Shear (12 hours), and Recovery (12 hours) were seeded at 5000 cells/mL on glass substrates and placed in an incubator for 6 hours to permit cell attachment and spreading. We used an inverted live cell fluorescence microscope (Leica DMI6000B, Leica Microsystems, Germany) with a 10X Phase Contrast objective to capture time-lapse images of cells every 15 minutes over 6 hours in a humid temperature-controlled PECON chamber maintained at 37°C and CO_₂_ levels at 5%. A motorized stage was used to capture multiple positions (N∼10) of each cell in the Control, Shear, and Recovery groups. Phase contrast time-lapse images were converted to 8-bit and drift-corrected using the Template Matching plugin in FIJI with bilinear interpolation^50^. Migrations of cells were tracked using a Manual Tracking Plugin after applying a centering correction. Cells that migrated a distance greater than three cell lengths were included in the analysis. Migration tracks were aligned to a common point of origin, and the overall displacement, mean, and instantaneous cell speeds were calculated using the (x,y) positions of cells at each time instant, t. Mean Square Displacement (MSD) and the directionality ratio (DR) were calculated using the DiPer code^51^ using overlapping time intervals. Based on this method, the MSD(n), representing the displacement for step size, n, is given by

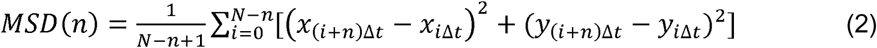

where N is the total number of displacements step size n and Δt the minimal time interval between points. Population averages of MSDs were plotted with time. The directionality ratio was calculated as

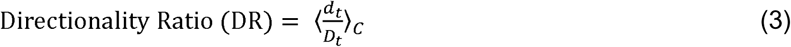

d _t_ represents the straight-line distance from the initial position of the cell to its position at time t, providing a direct measure of displacement from the starting point. D_t_ denotes the actual path length traveled by the cell from the start to its position at time t, which includes all turns and deviations. The brackets indicate an average of these values across C cells at time, t.

### RNA Isolation and RT-qPCR

Cells were washed with PBS, and TRIzol® reagent was added to each Petri dish for RNA extraction using the RNeasy Mini Kit (Qiagen). Two micrograms of RNA were used for cDNA synthesis with a high-capacity cDNA reverse transcription kit (Applied Biosystems). Quantitative real-time PCR (qRT-PCR) was conducted using PowerUp SYBR Green master mix with 10 ng of cDNA as the template. Fold changes were calculated using the 2^−ΔΔCT^ method and gene expression was normalized to GAPDH and no-shear controls. Data were analyzed across three biological replicates for each Control, Shear, and Recovery group in our study.

### Immunofluorescence

Cultured cells were rinsed twice with chilled PBS (Fisher Scientific), fixed with 4% paraformaldehyde for 15Lminutes, permeabilized with 0.1% Triton X-100 for 3Lminutes, and blocked using 5% bovine serum albumin (BSA) with 5% fetal bovine serum (FBS) for 1Lhour at room temperature for immunofluorescence staining. Specimens were incubated for 1 hour at room temperature with primary anti-vinculin antibody (1:500, Sigma Aldrich) followed by secondary Alexa Fluor 488-conjugated antibody (1:600, Molecular Probes, Invitrogen), with Alexa Fluor 568-phalloidin (1:200, Invitrogen) to stain F-actin. Samples were rinsed twice with PBS, nuclei were labeled with DAPI (1:500, Invitrogen) for 2Lminutes, rinsed, and mounted on coverslips with ProLong™ Gold Antifade Mountant (Invitrogen). Confocal images were acquired using a Leica SP5 confocal microscope. Images were analyzed using Fiji (ImageJ) to quantify cell spread area after aligning samples along the shear direction.

Stress fibers were quantified using confocal images of cells stained for actin, and a custom MATLAB code as described earlier^28^. This method involves segmenting the image to identify actin fibers using intensity thresholding and morphological operations. The lengths of individual stress fibers were calculated using the Hough transform of the segmented fibers from each image. Approximately 5-10 fibers were identified in each cell, and 15 cell images were analyzed for each group in the study.

### Statistical Analysis

Data from experiments performed in three replicates are reported as mean ± SD. Differences between groups were assessed using one-way analysis of variance (ANOVA) with Bonferroni post-hoc tests for individual comparisons. A p-value of < 0.05 was considered statistically significant. Non-significant differences are indicated as n.s.

